# Structure of SARS-CoV-2 M protein in lipid nanodiscs

**DOI:** 10.1101/2022.06.12.495841

**Authors:** Kimberly A. Dolan, Mandira Dutta, David M. Kern, Abhay Kotecha, Gregory A. Voth, Stephen G. Brohawn

## Abstract

SARS-CoV-2 encodes four structural proteins incorporated into virions, spike (S), envelope (E), nucleocapsid (N), and membrane (M). M plays an essential role in viral assembly by organizing other structural proteins through physical interactions and directing them to sites of viral budding. As the most abundant protein in the viral envelope and a target of patient antibodies, M is a compelling target for vaccines and therapeutics. Still, the structure of M and molecular basis for its role in virion formation are unknown. Here, we present the cryo-EM structure of SARS-CoV-2 M in lipid nanodiscs to 3.5 Å resolution. M forms a 50 kDa homodimer that is structurally related to the SARS-CoV-2 ORF3a viroporin, suggesting a shared ancestral origin. Structural comparisons reveal how intersubunit gaps create a small, enclosed pocket in M and large open cavity in ORF3a, consistent with a structural role and ion channel activity, respectively. M displays a strikingly electropositive cytosolic surface that may be important for interactions with N, S, and viral RNA. Molecular dynamics simulations show a high degree of structural rigidity and support a role for M homodimers in scaffolding viral assembly. Together, these results provide insight into roles for M in coronavirus assembly and structure.

## Introduction

Coronaviruses encode four structural proteins that are incorporated into mature enveloped virions: the transmembrane spike (S), membrane (M), and envelope (E) proteins and the soluble nucleocapsid (N) protein^1^. S proteins protrude from the virion, creating the eponymous corona in electron micrographs, and mediate fusion of viral and host cell membranes. E proteins form cationic viroporins that promote viral assembly and modulate the host immune response. N is an RNA-binding protein that packages the viral RNA genome. M organizes the assembly and structure of new virions and is essential for virus formation^2–5^. M is the most abundant membrane protein in the viral envelope and anti-M antibodies are found in plasma of patients infected with SARS-CoV-2 and other coronaviruses^6–11^. Based on its functional importance and immunogenicity, M has been proposed as a target for coronavirus vaccines or therapeutics.

In infected cells, M mediates virus assembly and budding by interacting with all other structural proteins and directing their localization to the ER-Golgi intermediate compartment (ERGIC)^1,12–19^. M is proposed to interact with E through its transmembrane region and S and N through a cytosolic C-terminal region^16–19^. ER export and Golgi localization sequences in M determine its subcellular localization and M, in turn, modulates localization and posttranslational processing of S to promote virion assembly^19,20^. Across a wide range of coronaviruses (including SARS-CoV-2, SARS-CoV-1, MERS, mouse hepatitis virus (MHV), infectious bronchitis virus, and transmittable gastroenteritis virus), M is required for minimal virus-like particle (VLP) formation in transfected cells^14,21–24^. M is insufficient for VLP formation alone, however, and co-required components vary in different systems. SARS-CoV-2 VLP formation requires M co-expression with S or N^23,24^.

M has further been implicated in modulating host antiviral innate immunity. M inhibits the innate immune response by interfering with MAVS-mediated signaling and interferon production^25,26^. In mouse models of infection, M expression results in lung epithelial cell apoptosis in vitro and in vivo and may contribute to lung injury and pulmonary edema found in severe disease^26^.

Despite its essential role in viral assembly and implication in pathogenesis, the molecular determinants of M function remain largely unknown. MHV M was proposed to adopt long and compact structures that differentially facilitate membrane bending and recruitment of other structural proteins based on low resolution tomographic analysis^15^. Intriguingly, a structural and evolutionary relationship between SARS-CoV-2 M and the accessory viroporin ORF3a was reported^27^ based on predicted homology to our experimental ORF3a structures^28^. The manner in which distinct functional roles for M and ORF3a can be achieved in the context of a shared architecture remains to be determined. Here, we report the cryo-EM structure of SARS-CoV-2 M in lipid nanodiscs and perform molecular dynamics simulations to provide insight into M structure, function, and dynamics.

## Results

We determined the structure of SARS-CoV-2 M in lipid nanodiscs. Full-length M was expressed in *Spodoptera frugiperda* (Sf9) cells with a cleavable C-terminal GFP tag. Gel filtration chromatography of protein extracted in DDM/CHS detergent shows M runs predominantly as a single species consistent with a 50 kDa homodimer. We do not observe evidence of specific higher order oligomerization at low concentrations by fluorescence size exclusion chromatography or at higher concentrations in large scale purifications (Fig. S1). SARS-CoV-2 ORF3a, in contrast, assembles into stable homodimers and homotetramers under similar conditions^28^.

We reconstituted homodimeric SARS-CoV-2 M in nanodiscs made from the scaffold protein MSP1E3D1 and lipids (DOPE:POPC:POPS in a 2:1:1 ratio) and determined its structure by cryo-EM (Figs. 1, S2, Table 1). The majority of M (189 of 222 amino acids per subunit) was de novo modeled in the cryo-EM map (Figs. 1, S2). The N-terminus (amino acids 1-16) and C-terminus (amino acids 205-222) are not resolved in the map and were not modeled. Loops connecting transmembrane helices (amino acids 36-42 and 71-78) are the least well resolved regions of the structure. The relatively weak density is consistent with a lack of stabilizing interactions between these and other M regions and likely indicates they adopt a range of conformations among particles used to generate the final map.

**Figure 1.**
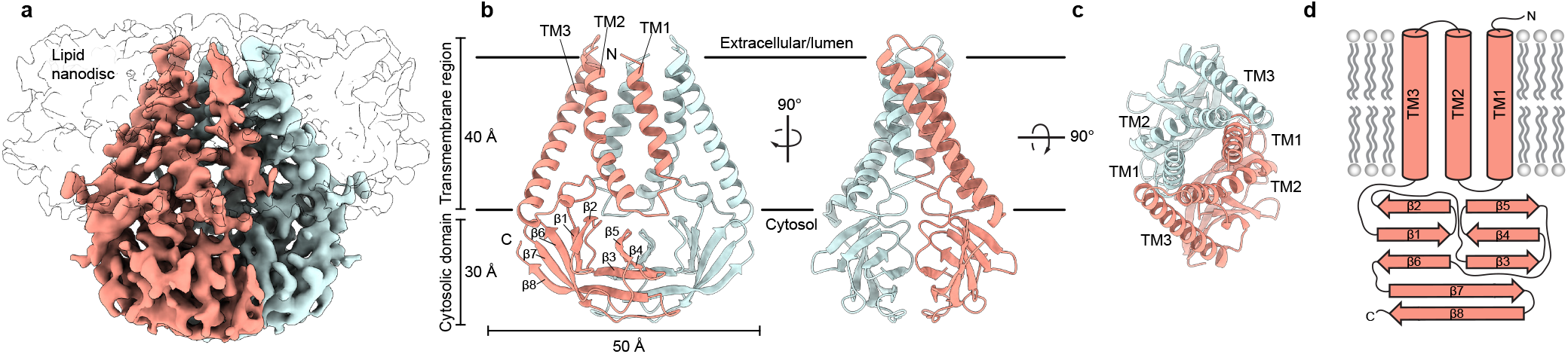
Structure of SARS-CoV-2 M in lipid nanodiscs. (a) 3.5 Å resolution cryo-EM map of SARS-CoV-2 M in MSP1E3D1 nanodiscs viewed from the membrane. One subunit is colored pink, and the second subunit is colored blue. Density corresponding to the lipid nanodisc is shown transparent. (b,c) Model of M viewed (b) from the membrane in two rotations and (c) from the extracellular or lumenal side. (d) Cartoon schematic of an M monomer with secondary structure elements indicated.

M is ~70 Å tall when viewed from the membrane with a ~40 Å transmembrane spanning region and ~30 Å cytosolic domain (CD) extending into the intracellular solution (Fig. 1). Each subunit contains an extracellular or lumenal N-terminus, three transmembrane helices (amino acids 17-36, 43-71, and 79-105) connected by short linkers, and a beta-strand rich C-terminal cytosolic domain. We note that AlphaFold and RoseTTAFold predicted M structures diverge substantially from the experimental structure (Fig. S3)^29,30^. In the predicted structures, TM1s are swapped between subunits in addition to differences in the relative positions of transmembrane and cytosolic domains.

Viewed from above, TM1-3 from each subunit is positioned along a flattened ellipse with a long major (~50 Å) and short minor (~16 Å) axis. Within each subunit, TM2-TM3 are closely juxtaposed and tightly packed while TM1-TM2 are more distant and loosely connected. The two subunits assemble along their long axis, with TM1 from one protomer forming extensive interactions with the TM2-TM3 unit of the second protomer. As they project towards the cytoplasm, the three transmembrane helices twist counterclockwise and splay outwards in the inner leaflet, creating an expanded ellipse with ~55 Å and ~40 Å axes at the intracellular leaflet.

The transmembrane region is connected to the cytosolic domain through a tight turn-helix-turn segment comprised of residues 106-116. Within the cytosolic domain, each protomer chain forms a pair of opposing β-sheets packed against one another in an eight stranded β-sandwich (Fig. 1B,D). The outer sheet is formed by strands β1, β2, β6, the N-terminal half of β7, and the C-terminal half of β8. The inner sheet is formed by strands β3, β4, β5, the C-terminal half of β7, and the N-terminal half of β8. The inner sheets from each protomer interact through a large (~690 Å^2^ buried surface area per chain) and complementary interface with residues L138, V139, V143, L145, F193, A195 contributing to a hydrophobic core surrounded by additional polar interactions.

Using Dali^31^ to compare the M structure to all experimentally determined protein structures returns SARS-CoV-2 ORF3a^28^ as the only structural homolog with a shared fold. Superposition of the two viral proteins reveals a similar fold topology and homodimeric assembly with an overall RMSD of 4.4 Å. Isolated transmembrane and cytosolic domains from individual protomers are better superimposed (RMSD = 2.7 Å and 2.5 Å, respectively) (Fig. 2). Substantial differences in M and ORF3a structure are observed in three regions: TM2-TM3, the transmembrane-cytosolic domain junction, and the cytoplasmic domain interface. TM1s of M and ORF3a are well superimposed, but TM2-TM3 of M are splayed further out into the membrane and are less twisted about the two-fold symmetry axis to create a flatter and tighter interaction surface. The angle between TM3 and the cytosolic domain is ~25° more acute in M. The cytosolic domains of M are rotated ~15° away from the symmetry axis relative to ORF3a, shifting the cytosolic domain interface between subunits further from the membrane.

**Figure 2.**
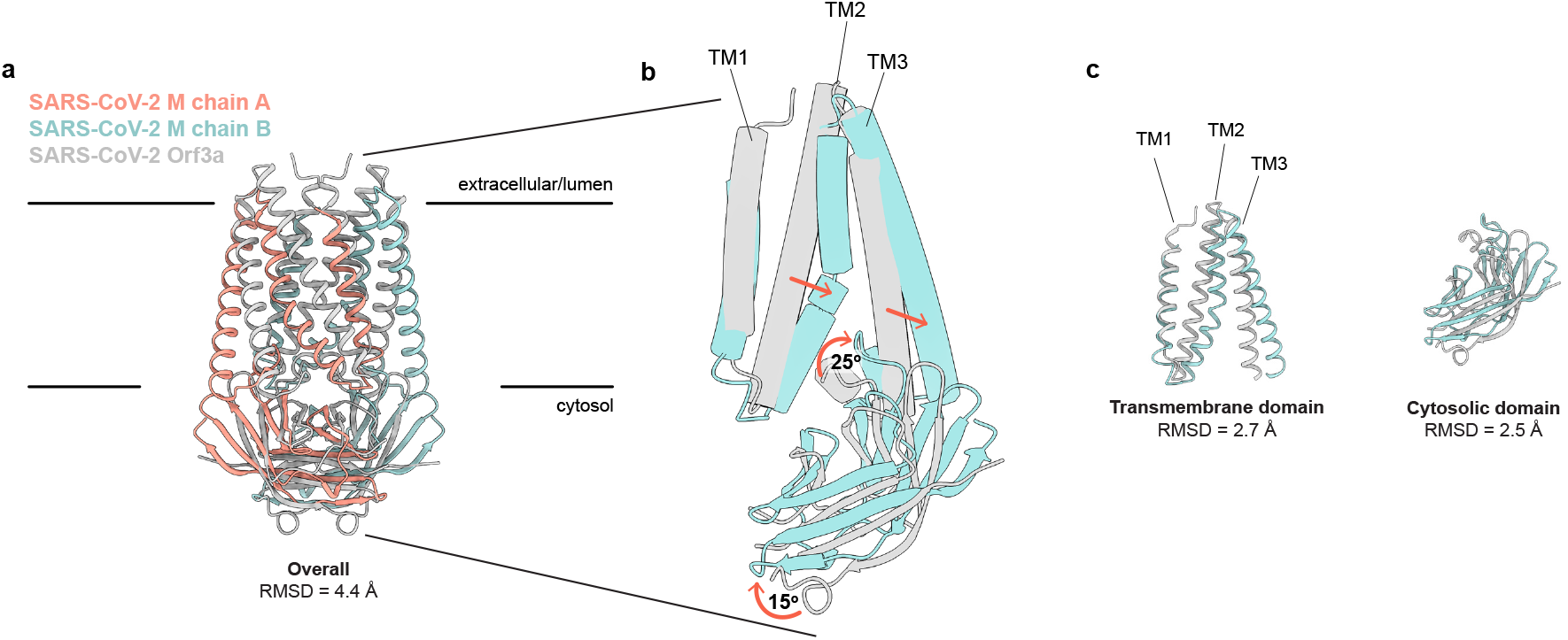
SARS-CoV-2 M and ORF3a proteins are structurally homologous. (a) Overlay of M and ORF3a structures. M is colored with one subunit pink and the second subunit blue and ORF3a is white. (b) Overlay of a single subunit indicating major conformational rearrangements. (c) Overlay of isolated transmembrane and cytosolic domains from each protein.

What are the consequences of the structural rearrangement in M relative to ORF3a? Association of subunits in the M homodimer creates a polar and, presumably, water-filled pocket, reminiscent of the polar cavity created between subunits in ORF3a. However, the included volumes are different in several key respects (Fig. 3). First, the M pocket is ~⅓by the number of close smaller with an enclosed volume of ~840 Å^3^ compared to ~1300 Å^3^ in ORF3a. Second, the M pocket is completely sealed by protein to the surrounding membrane and cytoplasm; no openings large enough for water passage are observed connecting the pocket and protein exterior. In contrast, ORF3a displays three pairs of tunnels connecting its internal cavity to the membrane and cytoplasm (two are displayed in Fig. 3). Third, the position of the M pocket and ORF3a cavity are different. In M, the gap between subunits is confined to the region between cytosolic domains because transmembrane helices from opposing subunits form tight interactions across the entire lipid bilayer. In ORF3a, the gap extends from the region between cytosolic domains to approximately halfway across the membrane because transmembrane helices are less tightly associated across the membrane inner leaflet.

**Figure 3.**
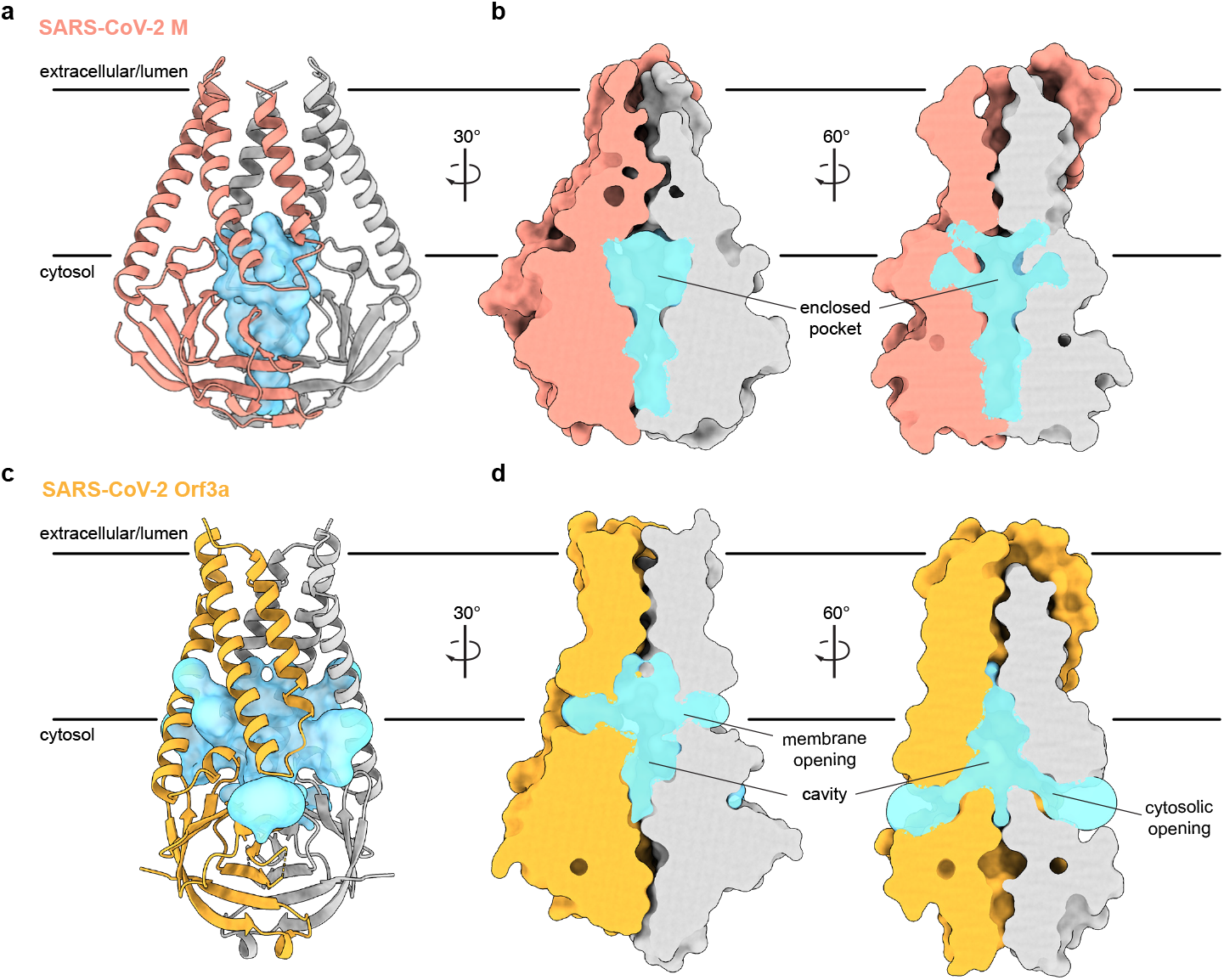
An enclosed polar pocket between cytosolic domains in M. (a) M shown as a cartoon and (b) surface with enclosed pocket volume calculated with CASTp^36^ shown as a blue surface. The enclosed pocket in M is formed between cytosolic domains and is sealed to the surrounding solution by protein. (c,d) Same as (a,b), but for SARS-CoV-2 ORF3a. The cavity in ORF3a begins closer to the lipid bilayer, extends approximately halfway across the membrane, and is open to surrounding solution and lipids through multiple openings^28^.

Another major difference in M and ORf3a structures is shown in Figure 4. The cytosolic domain of M is strikingly electropositive across nearly the entire exposed surface. Electropositive character is contributed by seventeen basic amino acids in three surface patches. The first covers the wide face of the cytosolic domains and consists of eight residues (R44, H125, R131, R146, H148, H155, R158, and R198). The second covers the narrow face of M and consists of four residues (R101, R105, R107, and R174). The third covers the underside of M and consists of five residues (K162, K166, K180, R186, and R200). ORF3a, in contrast, presents mixed electrostatic character with electropositive patches closer to the membrane and electronegative patches towards the cytoplasm. Such uniform electropositivity across the M cytosolic surface could facilitate the close juxtaposition of M present at high concentration in viral envelope with the negatively charged viral RNA genome.

**Figure 4.**
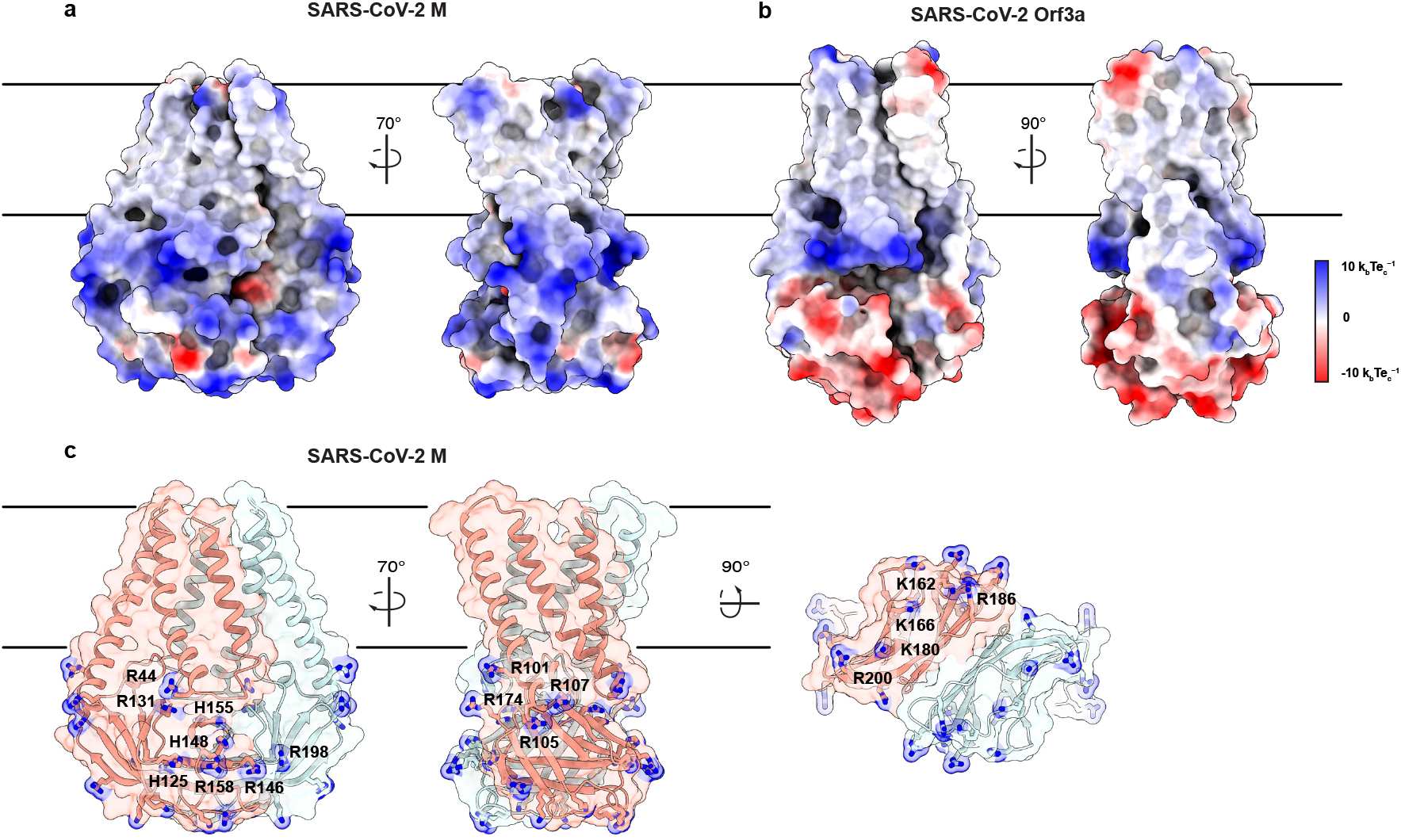
An electropositive cytosolic surface in M. (a,b) Views of the wide and narrow faces of (a) M and (b) ORF3a colored according to electrostatic surface potential from red (electronegative, −10 k_b_Te_c_^-1^) to blue (electropositive, +10 k_b_Te_c_^-1^). (c) Views of three electropositive surface patches on M cytosolic domains with basic residues labeled and shown as sticks with blue nitrogen atoms.

The large, complementary, and hydrophobic interface between transmembrane and cytosolic regions of subunits in the M structure suggests a structurally rigid core. However, a previous tomographic study of MHV suggesting that M adopts distinct long and compact structures^15^, M’s structural homology to the viroporin ORF3a^27,28^, and the dissociation of cytosolic regions shown in predicted SARS-CoV-2 M structures^29^ suggest the possibility that M is capable of undergoing large-scale structural rearrangements. Motivated by this discrepancy between the predicted dynamics of M and our experimental findings, we performed molecular dynamics (MD) simulations to gain insight into the potential for conformational changes in M.

We equilibrated M in a lipid environment and ran an all-atom MD simulation for 1.6 μs. Overall, we did not observe substantial conformational rearrangements in M during the simulation (Fig. 5A,B). Superposition of the experimental and final M structure following the simulation shows minor deviations through most of the protein (overall RMSD of 2.5 Å) (Fig. 5A). The largest difference is a shift in TM1 up towards the extracellular/lumenal side by approximately half a helical turn, enabled by rearrangement of the TM1-TM2 linker (Fig. 5A). This relatively subtle movement is consistent with weaker density for the TM1-TM2 linker in the cryo-EM map and fewer packing interactions for TM1 than TM2 or TM3. Per residue deviations ranged from ~ 1-4 Å and, aside from the movement of TM1, were similar between subunits and largest in the TM2-TM3 linker, transmembrane to cytosolic region connection, and loops connecting strands in the cytosolic domain. Minimal structural deviation was observed during the simulation within or between subunits as judged by the number of close Cα contacts, the angle between transmembrane and cytosolic regions, the distance between transmembrane regions, or the distance between cytosolic domains (Figs. 5D-H). Consistent with limited movement of the transmembrane region and a lack of evidence for lipid binding in the cryo-EM structure, no obvious enrichment of specific lipids around M was identified following simulation (Fig. S4). Finally, the internal M pocket remained similar in size and sealed from the surrounding solution throughout the simulation (Fig. 5I,J). We conclude that under these conditions M adopts a largely stable structure with minimal dynamic conformational rearrangement at physiological temperature.

**Figure 5.**
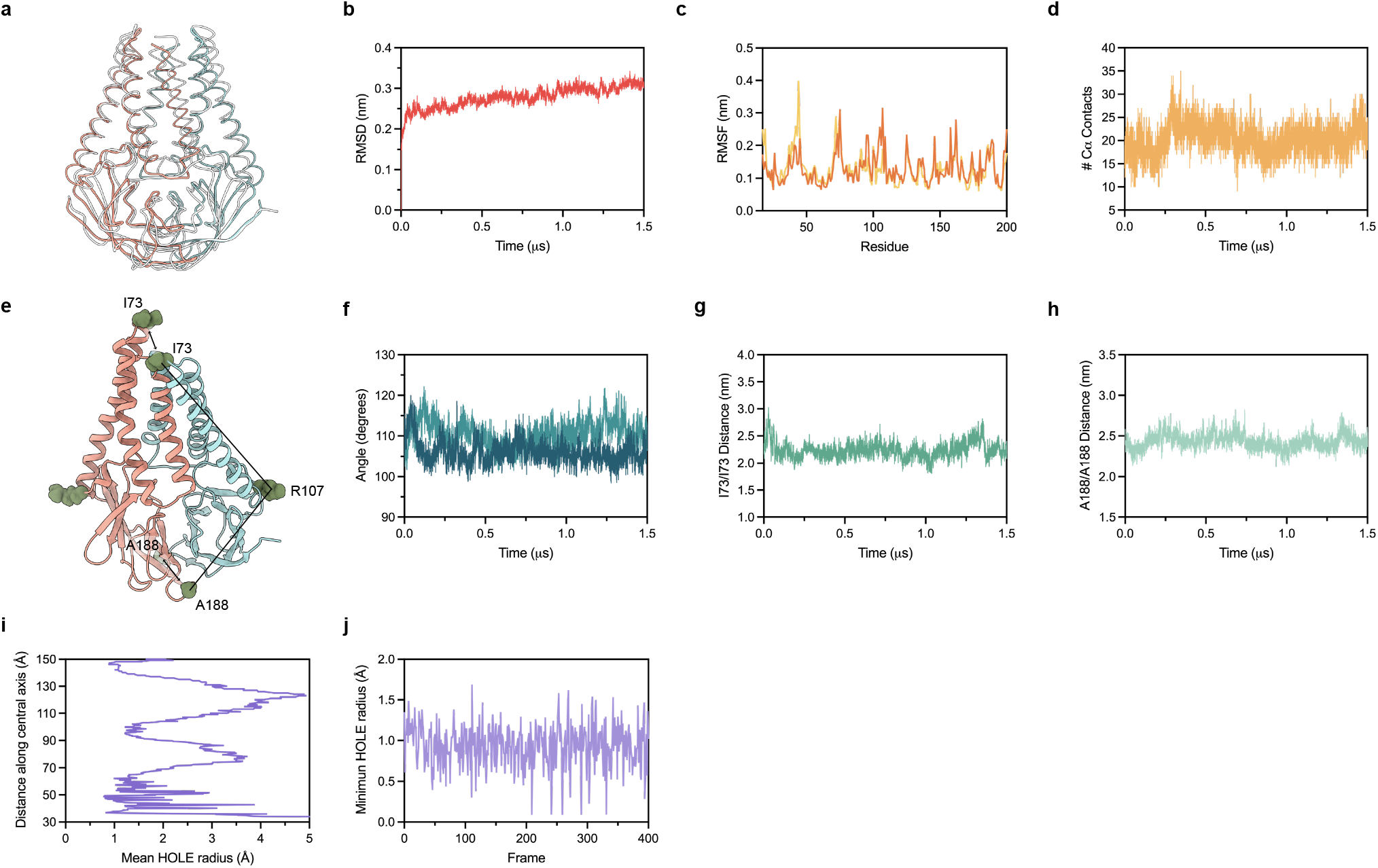
Molecular dynamics simulation of M. (a) Overlay of M cryo-EM structure (colored in pink and blue) and final structure (in white) following 1.6 μs all atom molecular dynamics simulation. (b) Overall RMSD between simulated and initial structure during simulation. (c) Root mean square fluctuation of protein residues in the simulation. Orange and yellow colors correspond to individual M protein chains. (d) Number of C-alpha contacts between two monomers. (e) Structural representation of distances and angles used for calculations in (f-h). (f) A188-R107-I73 angle plot for each monomer. One monomer has slightly higher values than the other. (g) Center of mass distance between I73 residues at the top of the TM2-TM3 linker. (h) Center of mass distance between A188 residues at the base of the cytosolic domains. (i) Mean radius of the enclosed pocket in M over the simulation trajectory versus distance along the symmetry axis. At its widest positions, the pocket is wide enough to accommodate two water molecules. (j) Minimum hole radius vs. the frame number in the simulation. The lack of substantial changes in radius indicates a stable pocket size and shape that does not open to solution during the simulation.

## Discussion

The structure of the SARS-CoV-2 M protein that we have obtained by cryo-EM reveals a homodimeric fold that is structurally homologous to the nonselective Ca^2+^ permeable cation channel of SARS-CoV-2, ORF3a. As with 3a, each subunit of M contains three transmembrane helices and a C-terminal beta sandwich domain.

However, the structure differs from ORF3a in several key ways that provides insight into how these structurally similar proteins can fill drastically different apparent roles in the coronavirus life cycle.

When viewed from the plane of the membrane, M is considerably wider and flatter than ORF3a, due to differences in transmembrane helix packing and a rotation about the central axis of the cytosolic domain. Among the consequences of this flattening out of M are distinct differences in the dimer interface across the membrane, where M shows a tighter dimer interface closer to the membrane outer leaflet as well as a gap between cytosolic domain subunits that forms an enclosed pocket lined by polar residues. In ORF3a, transmembrane regions are less closely opposed and a gap between subunits extends from halfway across the membrane to halfway down the cytosolic domains. The result is a larger cavity that is open to the membrane and cytoplasm. Mutations in the ORF3a cavity alter ion channel activity, consistent with the cavity forming part of the conduction path. Tight subunit association may therefore be important for the structural role of M, while loose subunit association that creates a large open cavity may be essential for the viroporin activity of ORF3a.

In further contrast to ORF3a, which was seen to form stable tetramers through electrostatic interactions between neighboring dimers, we see no evidence that M forms higher order oligomers under similar experimental conditions. Surface characteristics of the M dimer lend credence to the possibility that M exists solely as a dimer in the membrane–one striking feature of the M C-terminal beta sandwich domain is the presence of three sizable patches of positive charge that dominate its solvent exposed surface. Molecular dynamics simulations of M show that the dimer is stable and does not readily adopt alternate conformations at physiological temperature over the 1.6 μs trajectory. Taken together, these data suggest a purely structural role for M, whereby M mediates morphological changes in host cell membranes not through forming networks of M dimer-dimer interactions or through large-scale conformational changes, but rather through interactions with other SARS-CoV-2 structural proteins and perhaps negatively charged lipid headgroups or viral RNA.

M has also been shown to play a crucial role in viral assembly through protein-protein interactions with other coronavirus structural proteins such as N and S. Spike proteins are incorporated into coronavirus virions via interactions between the cytosolic tail of S and the cytosolic domain of M, however the precise details of this interaction are unknown^19^. In SARS-CoV-2, M and N or S are the minimal components required for forming VLPs when expressed heterologously in cells^23,24^. Several recent studies have suggested that the C-terminal domain of N is the site of interaction between SARS-CoV-2 M and N, but as with S a precise binding site has not been established^32,33^. It is possible that M and N interactions are mediated by favorable electrostatic interactions between negatively charged residues of the N CTD and one or more of the basic patches identified on the surface of the cytosolic domain of M. Through the sheer abundance of M dimers found in the membrane of SARS-CoV-2 virions, M and N together might facilitate VLP formation via a mechanism similar to the Gag precursor of HIV, where the high concentration of M C-terminal domains at the cytoplasmic membrane surface recruit and organize many N proteins that together physically extrude a membranous bud.

At present the World Health Organization puts the confirmed number of COVID-19 cases worldwide at nearly 530 million. Over the last two years the SARS-CoV-2 virus has undergone many mutations that have been extensively documented through sequencing efforts worldwide^34^. Despite this, the M protein sequence has remained virtually unchanged–a testament to the critical role that M plays in viral replication and assembly^35^. Furthermore, while only 20 amino acids in length, the N-terminus of M has been found to be highly immunogenic in COVID-19 patients^8–11^. M has also been shown to modulate innate immune response and could contribute to lung injury often seen in severe cases^25,26^. Given its clear importance in the coronavirus life cycle and pathogenicity, M presents an attractive target for therapeutics or vaccines. While M is well conserved across Coronaviridae (Fig. S5), it shows particularly high conservation between SARS-CoV-1 and SARS-CoV-2, with a sequence similarity of 90.54%, highlighting its potential as a therapeutic target for emergent coronaviruses in the future.

**Figure S1.**
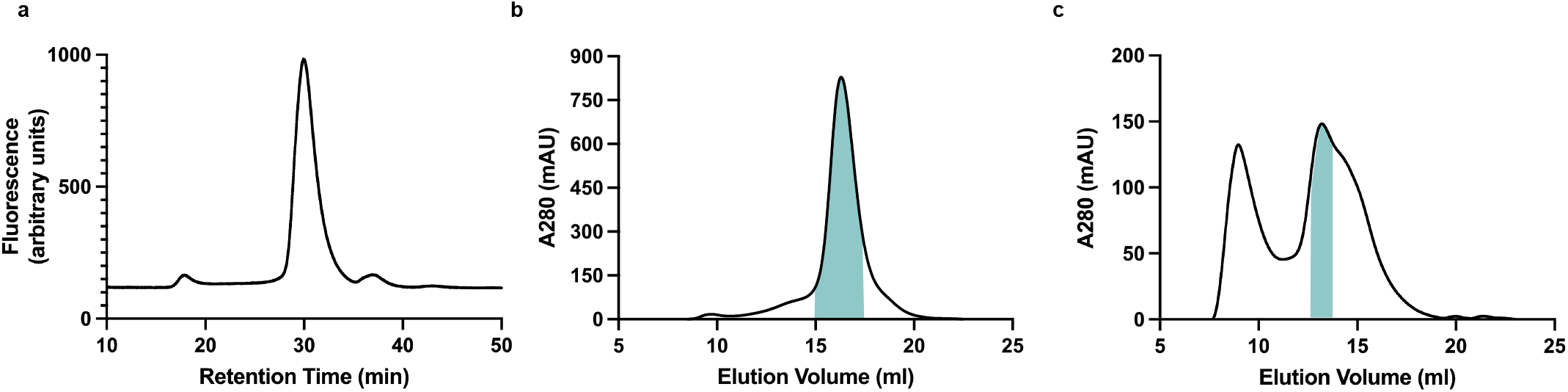
Purification and reconstitution of M. (a) Fluorescence size exclusion chromatogram of M expressed in insect cells and extracted in DDM/CHS. (b) Size exclusion chromatogram of M expressed in insect cells and extracted and purified in DDM/CHS. (c) Size exclusion chromatogram of M reconstituted into MSP1E3D1 lipid nanodiscs. Samples were run on Superose 6 columns. Blue bars indicate pooled fractions.

**Figure S2.**
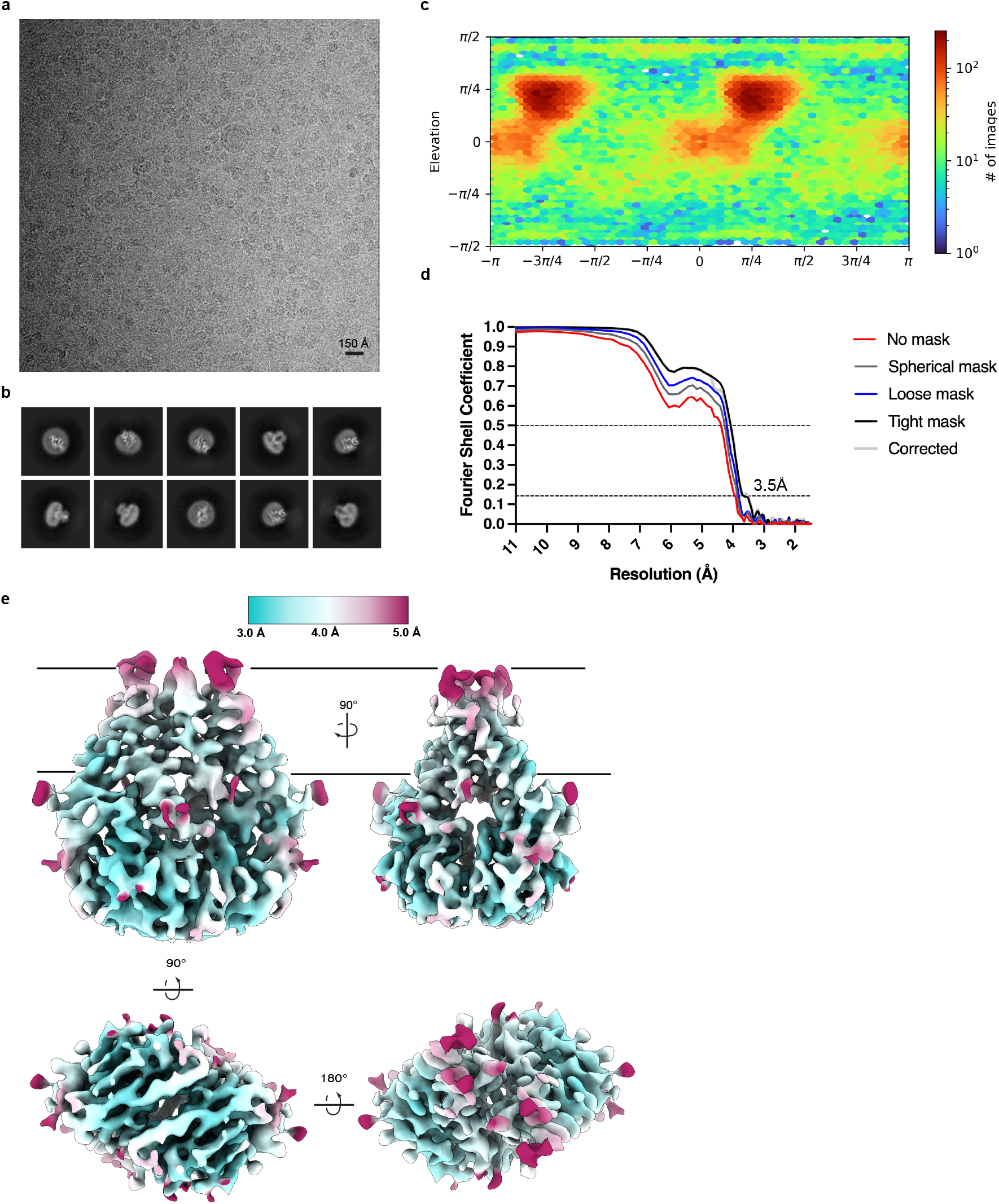
Cryo-EM processing and validation. (a) Representative micrograph, (b) selected 2D class averages, (c) angular distribution of particles used in final refinement, (d) Fourier shell correlation (FSC) relationships, and (e) local resolution estimated in CryoSPARC colored as indicated on the final map. Side, top, and bottom views are shown.

**Figure S3.**
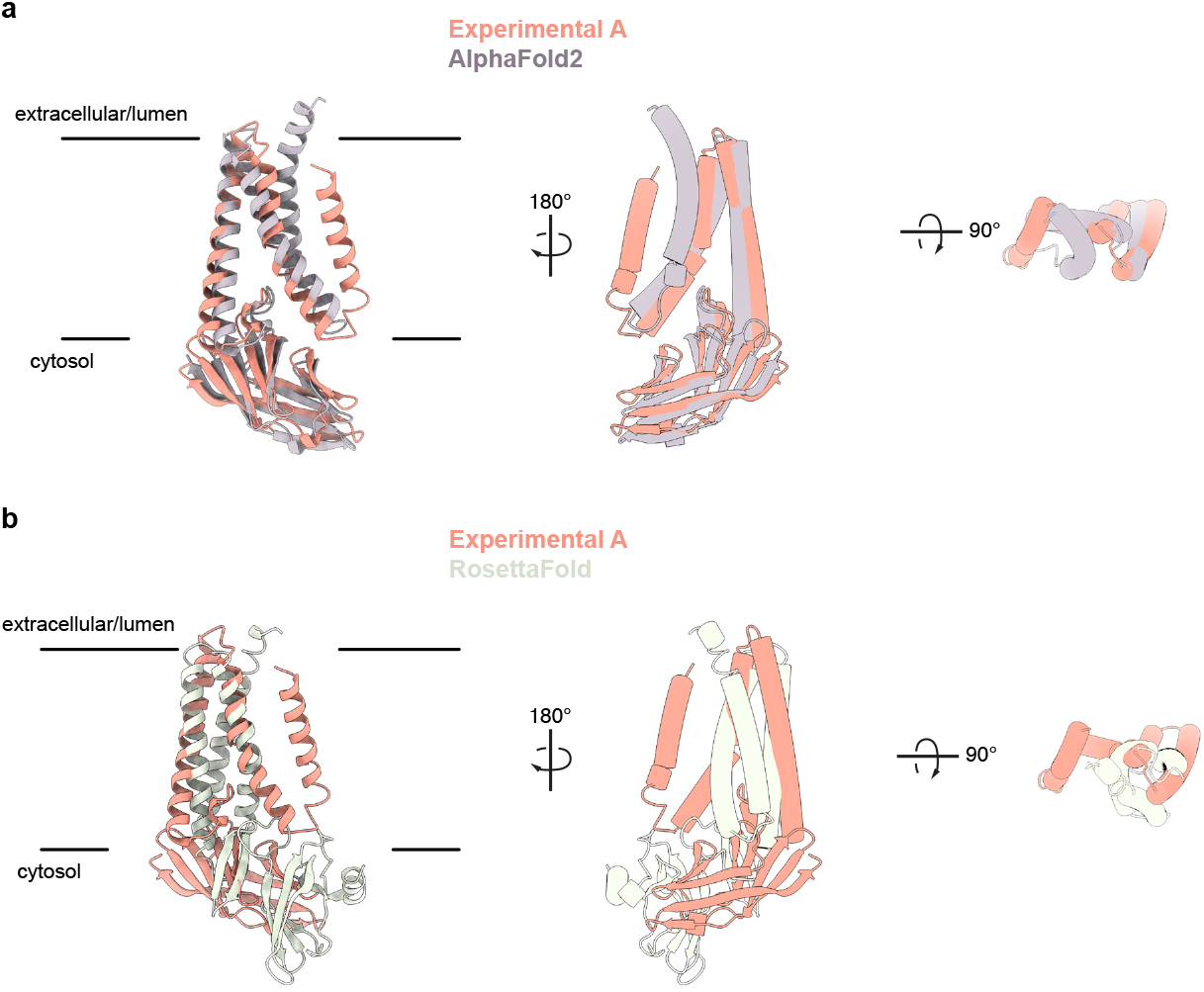
Comparison of experimentally determined and predicted M protein structures. (a) Overlay of the experimental and AlphaFold2 predicted M structures^30^. (b) Overlay of experimental and one RoseTTAfold predicted M structure^29^. Major differences are observed in chain topology in the transmembrane region and relative orientation of transmembrane and cytosolic domains.

**Figure S4.**
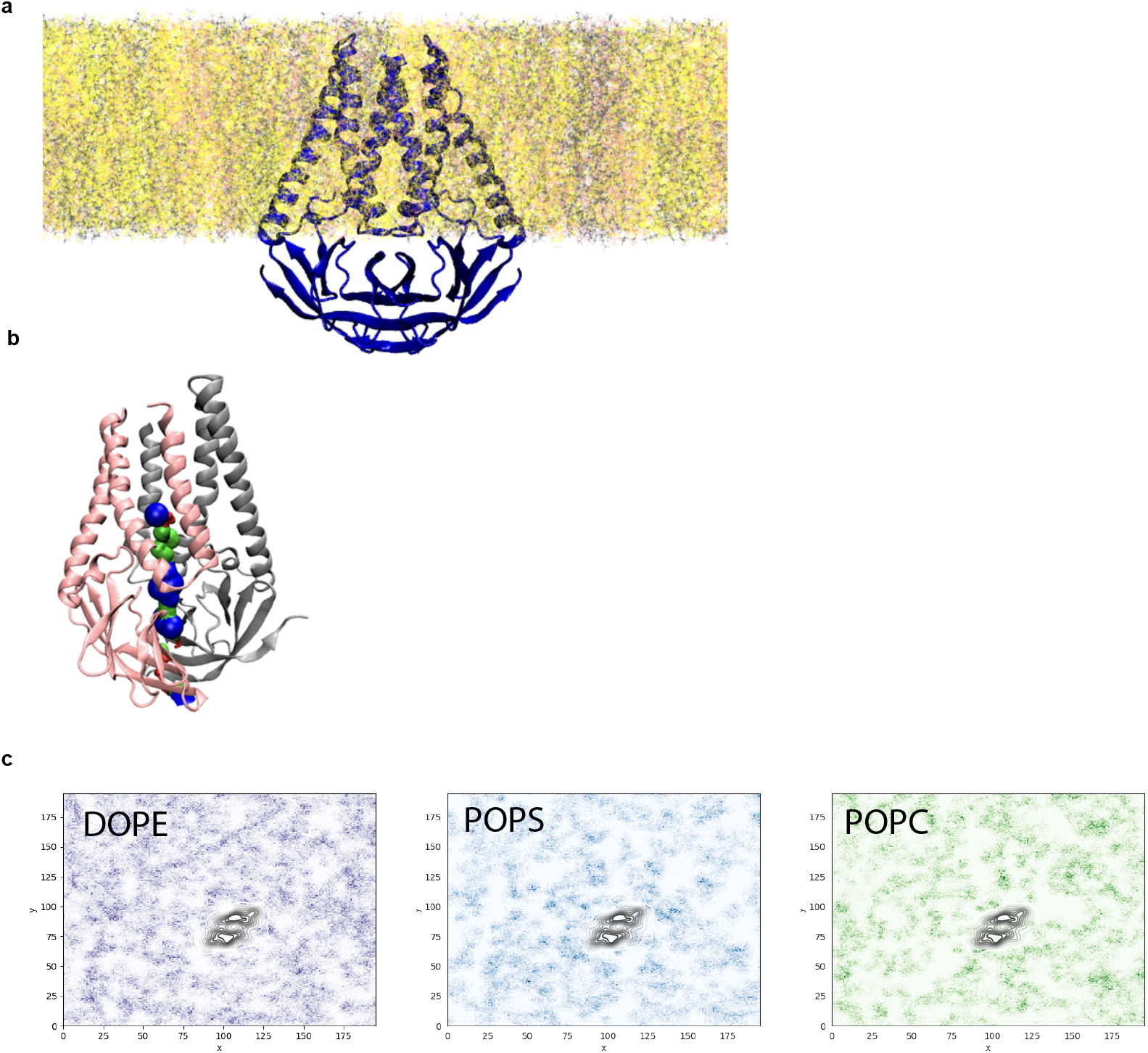
Analysis of pocket size and lipid interactions from molecular dynamics simulation. (a) Equilibrated M structure in lipid membrane consisting of DOPE:POPS:POPC in a 2:1:1 mass ratio. (b) Final M structure of the simulation shown with the pocket colored red where the radius is too small for a water molecule, green where it can accommodate a single water molecule, and blue where it can accommodate two water molecules. (c) Lipid sorting patterns. Darker regions indicate higher density of the lipids. Black color is the average location of the M-dimer. No obvious pattern of enrichment is observed.

**Figure S5.**
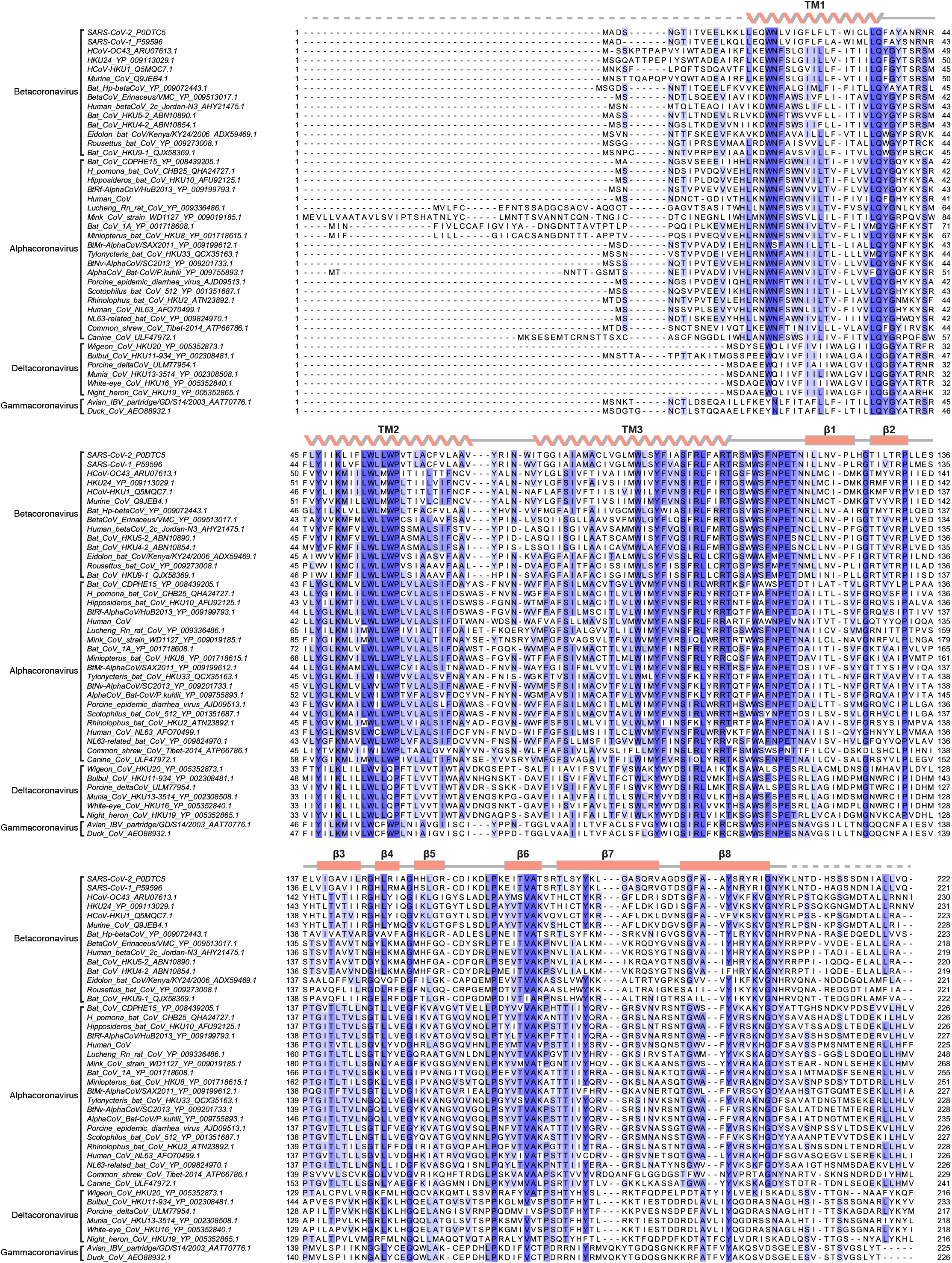
Sequence alignment of M proteins across *Coronaviridae*. (A) Alignment of forty-two M protein sequences colored by conservation in a ramp from white (not conserved) to dark blue (highly conserved). Accession numbers are indicated. Sequences were selected from representative species from each Coronavirus subgenus. Secondary structure from SARS-CoV-2 M is drawn above the alignment.

## Methods

### Cloning and protein expression

The coding sequence for SARS-Cov-2 M protein (Uniprot P0DTC5) was synthesized (IDT, Newark, NJ) and cloned into a vector based on the pACEBAC1 backbone (MultiBac; Geneva Biotech, Geneva, Switzerland) with an added C-terminal PreScission protease (PPX) cleavage site, linker sequence, superfolder GFP (sfGFP) and 7xHis tag, generating a construct for expression of M-SNS-LEVLFQGP-SRGGSGAAAGSGSGS-sfGFP-GSS-7xHis^37^. MultiBac cells were used to generate a Bacmid according to manufacturer’s instructions. Sf9 cells were cultured in ESF 921 medium (Expression Systems, Davis, CA) and P1 virus was generated from cells transfected with Escort IV reagent (MillaporeSigma, Burlington, MA) according to manufacturer’s instructions. P2 virus was then generated by infecting cells at 2 million cells/mL with P1 virus at a MOI ~0.1, with infection monitored by fluorescence and harvested at 72 hours. P3 virus was generated in a similar manner to expand the viral stock. The P2 or P3 viral stock was then used to infect Sf9 cells at 4 million cells/mL at a MOI ~2–5. At 72 hours, infected cells containing expressed M-sfGFP protein were harvested by centrifugation at 2500 x g for 10 minutes and frozen at −80°C.

### Protein purification

Infected Sf9 cells from 1 L of culture (~15 mL of cell pellet) were thawed in 100 mL of Lysis Buffer containing 50 mM HEPES, 150 mM KCl, 1mM EDTA pH 8. Protease inhibitors (Final Concentrations: E64 (1 μM), pepstatin A (1 μg/mL), soy trypsin inhibitor (10 μg/mL), benzamidine (1 mM), aprotinin (1 μg/mL), leupeptin (1 μg/mL), AEBSF (1 mM), and PMSF (1 mM)) were added to the lysis buffer immediately before use.

Benzonase (4 μl) was added after the cell pellet thawed. Cells were then lysed by sonication and centrifuged at 150,000 x g for 45 minutes. The supernatant was discarded, and residual nucleic acid was removed from the top of the membrane pellet using DPBS. Membrane pellets were scooped into a dounce homogenizer containing extraction buffer (50 mM HEPES, 150 mM KCl, 1 mM EDTA, 1% n-Dodecyl-β-D-Maltopyranoside (DDM, Anatrace, Maumee, OH), 0.2% cholesteryl hemisuccinate Tris salt (CHS, Anatrace, Maumee, OH) pH 8). A stock solution of 10% DDM, 2% CHS was dissolved and clarified by bath sonication in 200 mM HEPES pH 8 prior to addition to buffer to the indicated final concentration. Membrane pellets were then homogenized in extraction buffer and this mixture (150 mL final volume) was gently stirred at 4°C for 1.5 hours. The extraction mixture was centrifuged at 33,000 x g for 45 minutes and the supernatant, containing solubilized membrane protein, was bound to 4 mL of Sepharose resin coupled to anti-GFP nanobody for 1.5 hours at 4°C. The resin was then collected in a column and washed with 10 mL of buffer 1 (20 mM HEPES, 150 mM KCl, 1 mM EDTA, 0.025% DDM, 0.005% CHS, pH 7.4), 40 mL of buffer 2 (20 mM HEPES, 500 mM KCl, 1 mM EDTA, 0.025% DDM, 0.005% CHS, pH 7.4), and 10 mL of buffer 1. The resin was then resuspended in 6 mL of buffer 1 with 0.5 mg of PPX protease and rocked gently in the capped column for 2 hours. Cleaved M protein was then eluted with an additional 12 mL of wash buffer, spin concentrated to ~1 mL with Amicon Ultra spin concentrator 10 kDa cutoff (Millipore), and loaded onto a Superose 6 increase column (GE Healthcare, Chicago, IL) on an NGC system (Bio-Rad, Hercules, CA) equilibrated in buffer 1. Peak fractions containing M protein were then collected and spin concentrated prior to incorporation into nanodiscs.

### Nanodisc Formation

Freshly purified M protein in Buffer 1 was reconstituted into MSP1E3D1 nanodiscs with a mixture of lipids (DOPE:POPS:POPC at a 2:1:1 mass ratio, Avanti, Alabaster, Alabama) at a final molar ratio of 1:4:400 ( M:MSP1E3D1:lipid).

20 mM solubilized lipid in lipid dilution buffer (20 mM HEPES, 150 mM KCl, pH 7.4) was mixed with additional DDM/CHS detergent and M protein at 4°C for 30 minutes before addition of purified MSP1E3D1. This addition brought the final concentrations to approximately 10 μM M protein, 40 μM MSP1E3D1,4 mM lipid mix,10 mM DDM, and 1.7 mM CHS. The solution with MSP1E3D1 was mixed at 4°C for 15 minutes before addition of 150 mg of Biobeads SM2. Biobeads (washed into methanol, water, and then Nanodisc Formation Buffer) were weighed after liquid was removed by pipetting (damp weight). This final mixture was then gently tumbled at 4°C overnight (~ 12 hours). Supernatant was cleared of beads by letting large beads settle and carefully removing liquid with a pipette. Sample was spun for 10 minutes at 21,000 x g before loading onto a Superose 6 increase column in 20 mM HEPES, 150 mM KCl, pH 7.4. Peak fractions corresponding to M protein in MSP1E3D1 were collected, 10 kDa cutoff spin concentrated and used for grid preparation. MSP1E3D1 was prepared as previously described^38^ without cleavage of the His-tag.

### Cryo-EM sample preparation and data collection

M in MSP1E3D1 was prepared at a final concentration of 1.3 mg/mL. Concentrated sample was cleared by a 10-minute 21,000 x g spin at 4°C prior to grid preparation. 3.4 μl of protein was applied to freshly glow discharged Holey Carbon, 300 mesh R 1.2/1.3 gold grids (Quantifoil, Großlöbichau, Germany) and plunge frozen in liquid ethane using a FEI Vitrobot Mark IV (ThermoFisher Scientific) was used with 4°C, 100% humidity, 1 blot force, a wait time of ~5 seconds, and a 3 second blot time.

Grids were clipped and sent to Thermo Fisher Scientific RnD division in Eindhoven, The Netherlands for data collection. Grids were loaded onto a Krios G4 microscope equipped with a Cold Field Emission gun (CFEG) and operated at 300 kV. Data were collected on a Falcon 4 detector mounted behind a Selectris X energy filter. The slit width of the energy filter was set to 10eV. 7,588 movie stacks containing 1251 frames were collected with EER (electron event representation) mode^39^ of Falcon 4 detector at a magnification of 165,000 corresponding to a pixel size of 0.727 Å. Each movie stack was recorded with a total dose of 50e-/Å2 on sample and a defocus range between 0.5 to 1.2 μm.

See Table 1 for detailed data collection statistics.

### Cryo-EM data processing

Motion correction and dose weighting were performed on all 7,588 videos using RELION 4.0’s implementation of MotionCor2 at 0.727 Å per pixel. Contrast transfer function (CTF) parameters were fit with CTFFIND-4.1. Template-free auto-picking of particles was performed with RELION 4.0’s Laplacian-of-Gaussian filter on movies CTF fit to 5.0 Å or better, yielding an initial set of 2,379,507 particles. These particles were then extracted at a 288-pixel box size and transferred to cryoSPARC v.3.2 for two-dimensional classification.

Iterative rounds of 2D classification resulted in a set of 31,126 particles which were then extracted in RELION 4.0 and their coordinates were used in the Topaz particle-picking pipeline^40^. Topaz training, picking, and extraction yielded 2,376,190 particles which were then subjected to one round of 2D classification in RELION 4.0 to remove obvious noise. The resulting 2,007,561 particles were extracted and then iteratively 2D classified in cryoSPARC v3.2, resulting in a set of 54,747 ‘good’ particles.

Both the initial auto-picked particle set and subsequent Topaz particle set were lacking in good 2D classes of side views, so a subset of 13,698 particles from the best side view classes were extracted in RELION 4.0 and used to train and pick new particles in Topaz. As before, the resulting 2,186,648 particles were subjected to one round of 2D classification in RELION 4.0 then imported into cryoSPARC v3.2 for further 2D classification.

Good particles from the initial auto-picked particle set and both Topaz particle sets were pooled and duplicates within 100 Å were removed to yield 105,535 particles. These particles were extracted and imported into cryoSPARC v3.2 for 3 rounds of 2D classification to remove remaining junk. An ab initio reconstruction of the remaining 69,182 particles was performed to provide an initial volume and a subsequent non-uniform refinement (C2, 2 extra passes, 16 Å initial resolution) produced a map with a 4.0 Å overall resolution. This map was post-processed in RELION 4.0 and used for Bayesian particle polishing.

The resulting ‘shiny’ particles were imported back into cryoSPARC v3.2 for one additional round of 2D classification. The final 64,966 particles were used to generate a new ab initio and a subsequent non-uniform refinement (C2, 2 extra passes, 16 Å initial resolution, 1.5 adaptive window factor) yielded the final map at 3.5 Â nominal resolution.

### Modeling, Refinement, and Analysis

cryoSPARC sharpened cryo-EM maps were used to de novo model M using Coot^41^. The model was real space refined in Phenix^42^ and validated using Molprobity^43^. Cavity and tunnel measurements were made with CASTp^36^. Comparisons to the structure database were performed with DALI^31^. Figures were prepared using ChimeraX^44^, Prism, Adobe Photoshop, and Adobe Illustrator software.

### Fluorescence Size Exclusion Chromatography (FSEC)

Sf9 cells (~4 million) from the third day of infection were pelleted, frozen, and then thawed into extraction buffer (20mM Tris pH 8, 150 mM KCl, all protease inhibitors used for protein purification, 1 mM EDTA, 1% DDM). Extraction was performed at 4°C for 1 hour and lysate was then pelleted at 21,000 x g at 4°C for 1 hour to clear the supernatant. Supernatant was then run on a Superose 6 Increase column with fluorescence detection for GFP into 20 mM HEPES pH 7.4, 150 mM KCl, 0.025% DDM, 0.005% CHS.

### Molecular dynamics

The initial MD system of M-protein and lipid bilayer was built using CHARMM-GUI Membrane Builder^45–48^. A 20×20 nm lipid bilayer membrane was taken with a mixture of DOPE, POPS and POPC lipids in a 2:1:1 mass ratio. A fully hydrated bilayer was built around the M protein, centering the transmembrane region close to the lipid bilayer center. To neutralize the system, a 0.15M KCl salt concentration was used. The simulations were performed on GROMACS MD simulation package^49^ with the CHARMM36m force-field^50^. An initial minimization of the system was carried out following six-steps protocols provided on CHARMM-GUI^51^. A time step of 2fs was used with periodic boundary conditions for the simulations. A simulation temperature of 310.15 K was maintained with a Nose-Hoover thermostat^52,53^ and a coupling time constant of 1.0 ps in GROMACS. The pressure was set at 1bar with a Berendsen barostat^54^ during initial relaxation. For the production runs, the Parrinello-Rahman barostat was used semi-isotropically with the compressibility of 4.5 x 10^-5^ and a coupling time constant of 5.0 ps^55,56^. For the non-bonded interactions a switching function between 1.0 and 1.2 nm was used. The long-range electrostatics were computed using Particle Mesh Ewald (PME)^57^. The LINCS algorithm was used to constrain hydrogen bonds^58^. We performed 1.6 μs production run for the system and used Frontera (TACC), and Midway2 (Research Computing Center at the University of Chicago) to run these simulations.

The RMSD of the protein and RMSF per residue (Fig. 5B, C) were calculated using the GROMACS module. The center-of-mass (COM) distances between two residues (Fig. 5G, H), number of Cα contacts between two monomers (Fig. 5D), and angles between transmembrane and cytosolic regions (Fig. 5F) were also calculated using the GROMACS package^49^. The analysis of the M pocket was performed using the HOLE program^59^ implemented in MDAnalysis^60^ (Fig. 5I, J). In Fig. 5J, each frame was taken at a 4ns time step. The lipid distribution around the M protein was calculated using the MDAnalysis Python packages^60^ (Fig. S4). Visual Molecular Dynamics (VMD) and PyMOL were used as visualization software.

## Data and reagent availability

All data and reagents associated with this study are publicly available. The final model is in the PDB under 8CTK, the final map is in the EMDB under EMD-26993, and micrographs (original and motion corrected) and final particle stack are deposited in EMPIAR.

## Acknowledgements

We thank Dan Toso, Jonathan Remis, and Paul Tobias for support collecting preliminary data at the Cal-Cryo EM facility. We thank Thermo Fisher Scientific for microscope access. We thank Diana Bautista and members in Brohawn and Bautista labs for feedback on the project. We thank Savitha Sridharan and Hillel Adesnik for the initial M DNA. SGB is a New York Stem Cell Foundation-Robertson Neuroscience Investigator. This work was funded in part by the New York Stem Cell Foundation, a Sloan Research Fellowship (to SGB), a Fast Grants Award from Emergent Ventures at the Mercatus Center, George Mason University (to SGB, Diana Bautista, and Hillel Adesnik at UC Berkeley), an NSF Graduate Research Fellowship (to KD), and an NSF RAPID grant CHE-2029092 (MD and GAV). Computer simulations were carried out on the Frontera supercomputer at the Texas Advanced Computer Center (TACC) as funded by the National Science Foundation (OAC-1818253), as well as on the Midway2 cluster at the Research Computing Center (RCC) of the University of Chicago.

## Contributions

K.A.D., D.M.K., and S.G.B. conceived of the project. D.M.K. generated initial constructs and performed expression testing. K.A.D. generated final constructs and performed expression, purification, reconstitution, cryo-EM sample preparation, and cryo-EM data processing. A.K. collected cryo-EM data at Thermo Fisher Scientific. K.D. and S.G.B. built and refined the atomic model. M.D. and G.A.V. designed, performed, and analyzed molecular dynamics simulations. G.A.V. and S.G.B. secured funding and supervised research. K.A.D. and S.G.B. wrote the manuscript with input from all authors.

---

**Table S1.** Cryo-EM data collection, processing, refinement, and modeling data.

| Data collection | SARS-CoV-2 M |
| --- | --- |
| PDB | 8CTK |
| EMDB | 26993 |
| Total movies | 7588 |
| Magnification | 165,000 x |
| Voltage (KV) | 300 |
| Electron exposure (e<sup>-</sup>/Å<sup>2</sup>) | 50 |
| Defocus range (um) | -0.5 to -1.2 |
| Super resolution pixel size (Å<sup>2</sup>) | 0.3635 |
| Binned pixel size (Å<sup>2</sup>) | 0.727 |
| **Processing** |  |
| Initial particle images (no.) | 2,007,561 |
| Final particle images (no.) | 64,966 |
| Map resolution Masked (Å, FSC = 0.143) | 3.52 |
| Symmetry imposed | C2 |
| **Refinement** |  |
| Model resolution (Å, FSC = 0.143 / FSC = 0.5) | 3.2 / 3.5 |
| Map-sharpening B factor (Å<sup>2</sup>) | -145 |
| Composition |  |
| Number of atoms | 3042 |
| Number of protein residues | 376 |
| Number of ligands | 0 |
| RMS deviations |  |
| Bond lengths (Å) | 0.003 |
| Bond angles (Å) | 0.690 |
| Validation |  |
| MolProbity score | 1.47 |
| Clashscore | 4.37 |
| Ramachandran plot |  |
| Favored (%) | 96.24 |
| Allowed (%) | 3.76 |
| Disallowed (%) | 0 |
| Rotamer outliers (%) | 0 |
| Mean B factor (Å<sup>2</sup>) |  |
| Protein | 42.42 |

## References

1. Masters, P. S. The Molecular Biology of Coronaviruses. Adv Virus Res 66, 193–292 (2006).

2. Sturman, L. S., Holmes, K. V. & Behnke, J. Isolation of coronavirus envelope glycoproteins and interaction with the viral nucleocapsid. J Virol 33, 449–462 (1980).

3. Armstrong, J., Niemann, H., Smeekens, S., Rottier, P. & Warren, G. Sequence and topology of a model intracellular membrane protein, E1 glycoprotein, from a coronavirus. Nature 308, 751–752 (1984).

4. Yu, A. et al. A multiscale coarse-grained model of the SARS-CoV-2 virion. Biophys J 120, 1097–1104 (2021).

5. Finkel, Y. et al. The coding capacity of SARS-CoV-2. Nature 589, 125–130 (2021).

6. Godet, M., L’Haridon, R., Vautherot, J.-F. & Laude, H. TGEV corona virus ORF4 encodes a membrane protein that is incorporated into virions. Virology 188, 666–675 (1992).

7. He, Y., Zhou, Y., Siddiqui, P., Niu, J. & Jiang, S. Identification of Immunodominant Epitopes on the Membrane Protein of the Severe Acute Respiratory Syndrome-Associated Coronavirus. J Clin Microbiol 43, 3718–3726 (2005).

8. Hotop, S.-K. et al. Peptide microarrays coupled to machine learning reveal individual epitopes from human antibody responses with neutralizing capabilities against SARS-CoV-2. Emerg Microbes Infec 11, 1037–1048 (2022).

9. Heffron, A. S. et al. The landscape of antibody binding in SARS-CoV-2 infection. Plos Biol 19, e3001265 (2021).

10. Martin, S. et al. SARS-CoV-2 integral membrane proteins shape the serological responses of COVID-19 patients. Iscience 24, 103185 (2021).

11. Jörrißen, P. et al. Antibody Response to SARS-CoV-2 Membrane Protein in Patients of the Acute and Convalescent Phase of COVID-19. Front Immunol 12, 679841 (2021).

12. Haan, C. A. M. de, Smeets, M., Vernooij, F., Vennema, H. & Rottier, P. J. M. Mapping of the Coronavirus Membrane Protein Domains Involved in Interaction with the Spike Protein. J Virol 73, 7441–7452 (1999).

13. Cavanagh, D. Coronaviridae: a review of coronaviruses and toroviruses. Coronaviruses Special Emphas First Insights Concern Sars 1–54 (2005) doi:10.1007/3-7643-7339-3_1.

14. Siu, Y. L. et al. The M, E, and N Structural Proteins of the Severe Acute Respiratory Syndrome Coronavirus Are Required for Efficient Assembly, Trafficking, and Release of Virus-Like Particles. J Virol 82, 11318–11330 (2008).

15. Neuman, B. W. et al. A structural analysis of M protein in coronavirus assembly and morphology. J Struct Biol 174, 11–22 (2011).

16. Kuo, L., Hurst-Hess, K. R., Koetzner, C. A. & Masters, P. S. Analyses of Coronavirus Assembly Interactions with Interspecies Membrane and Nucleocapsid Protein Chimeras. J Virol 90, 4357–4368 (2016).

17. Kuo, L., Koetzner, C. A. & Masters, P. S. A key role for the carboxy-terminal tail of the murine coronavirus nucleocapsid protein in coordination of genome packaging. Virology 494, 100–107 (2016).

18. Lim, K. P. & Liu, D. X. The Missing Link in Coronavirus Assembly: Retention Of The Avian Coronavirus Infectious Bronchitis Virus Envelope Protein In The Pre-Golgi Compartments And Physical Interaction Between The Envelope And Membrane Proteins. J Biol Chem 276, 17515–17523 (2001).

19. Boson, B. et al. The SARS-CoV-2 envelope and membrane proteins modulate maturation and retention of the spike protein, allowing assembly of virus-like particles. J Biological Chem 296, 100111 (2020).

20. Perrier, A. et al. The C-terminal domain of the MERS coronavirus M protein contains a trans-Golgi network localization signal. J Biological Chem 294, 14406–14421 (2019).

21. Vennema, H. et al. Nucleocapsid-independent assembly of coronavirus-like particles by co-expression of viral envelope protein genes. Embo J 15, 2020–2028 (1996).

22. Hsieh, P.-K. et al. Assembly of Severe Acute Respiratory Syndrome Coronavirus RNA Packaging Signal into Virus-Like Particles Is Nucleocapsid Dependent. J Virol 79, 13848–13855 (2005).

23. Xu, R., Shi, M., Li, J., Song, P. & Li, N. Construction of SARS-CoV-2 Virus-Like Particles by Mammalian Expression System. Frontiers Bioeng Biotechnology 8, 862 (2020).

24. Plescia, C. B. et al. SARS-CoV-2 viral budding and entry can be modeled using BSL-2 level virus-like particles. J Biological Chem 296, 100103 (2020).

25. Fu, Y.-Z. et al. SARS-CoV-2 membrane glycoprotein M antagonizes the MAVS-mediated innate antiviral response. Cell Mol Immunol 18, 613–620 (2021).

26. Sui, L. et al. SARS-CoV-2 Membrane Protein Inhibits Type I Interferon Production Through Ubiquitin-Mediated Degradation of TBK1. Front Immunol 12, 662989 (2021).

27. Tan, Y. et al. Unification and extensive diversification of M/Orf3-related ion channel proteins in coronaviruses and other nidoviruses. Virus Evol 7, veab014 (2021).

28. Kern, D. M. et al. Cryo-EM structure of SARS-CoV-2 ORF3a in lipid nanodiscs. Nat Struct Mol Biol 573–582 (2021) doi:10.1038/s41594-021-00619-0.

29. Heo, L. & Feig, M. Modeling of Severe Acute Respiratory Syndrome Coronavirus 2 (SARS-CoV-2) Proteins by Machine Learning and Physics-Based Refinement. Biorxiv 2020.03.25.008904 (2020) doi:10.1101/2020.03.25.008904.

30. Jumper, J. et al. Highly accurate protein structure prediction with AlphaFold. Nature 1–11 (2021) doi:10.1038/s41586-021-03819-2.

31. Holm, L. Structural Bioinformatics, Methods and Protocols. Methods Mol Biology 2112, 29–42 (2020).

32. Cubuk, J. et al. The SARS-CoV-2 nucleocapsid protein is dynamic, disordered, and phase separates with RNA. Nat Commun 12, 1936 (2021).

33. Lu, S. et al. The SARS-CoV-2 nucleocapsid phosphoprotein forms mutually exclusive condensates with RNA and the membrane-associated M protein. Nat Commun 12, 502 (2021).

34. Hadfield, J. et al. Nextstrain: real-time tracking of pathogen evolution. Bioinformatics 34, 4121–4123 (2018).

35. Cagliani, R., Forni, D., Clerici, M. & Sironi, M. Computational Inference of Selection Underlying the Evolution of the Novel Coronavirus, Severe Acute Respiratory Syndrome Coronavirus 2. J Virol 94, e00411–20 (2020).

36. Tian, W., Chen, C., Lei, X., Zhao, J. & Liang, J. CASTp 3.0: computed atlas of surface topography of proteins. Nucleic Acids Res 46, W363–W367 (2018).

37. Kern, D. M. & Brohawn, S. G. SARS-CoV-2 3a expression, purification, and reconstitution into lipid nanodiscs. Methods Enzymol 653, 207–235 (2021).

38. Ritchie, T. K. et al. Chapter 11 - Reconstitution of membrane proteins in phospholipid bilayer nanodiscs. Methods in enzymology 464, 211–231 (2009).

39. Guo, H. et al. Electron-event representation data enable efficient cryoEM file storage with full preservation of spatial and temporal resolution. Iucrj 7, 860–869 (2020).

40. Bepler, T. et al. Positive-unlabeled convolutional neural networks for particle picking in cryo-electron micrographs. Nature Methods 16, 1153–1160 (2019).

41. Emsley, P., Lohkamp, B., Scott, W. G. & Cowtan, K. Features and development of Coot. Acta crystallographica Section D, Biological crystallography 66, 486–501 (2010).

42. Afonine, P. V. et al. Real-space refinement in PHENIX for cryo-EM and crystallography. Acta crystallographica. Section D, Structural biology 74, 531–544 (2018).

43. Williams, C. J. et al. MolProbity: More and better reference data for improved all-atom structure validation. Protein science: a publication of the Protein Society 27, 293–315 (2018).

44. Goddard, T. D. et al. UCSF ChimeraX: Meeting modern challenges in visualization and analysis. Protein science: a publication of the Protein Society 27, 14–25 (2018).

45. Jo, S., Kim, T., Iyer, V. G. & Im, W. ChArMM-GUI: A web-based graphical user interface for CHARMM. J Comput Chem 29, 1859–1865 (2008).

46. Brooks, B. R. et al. CHARMM: The biomolecular simulation program. J Comput Chem 30, 1545–1614 (2009).

47. Lee, J. et al. CHARMM-GUI Input Generator for NAMD, GROMACS, AMBER, OpenMM, and CHARMM/OpenMM Simulations Using the CHARMM36 Additive Force Field. J Chem Theory Comput 12, 405–413 (2016).

48. Jo, S., Kim, T. & Im, W. Automated Builder and Database of Protein/Membrane Complexes for Molecular Dynamics Simulations. PLoS ONE 2, e880 (2007).

49. Abraham, M. J. et al. GROMACS: High performance molecular simulations through multi-level parallelism from laptops to supercomputers. Softwarex 1, 19–25 (2015).

50. Huang, J. et al. CHARMM36m: an improved force field for folded and intrinsically disordered proteins. Nat Methods 14, 71–73 (2017).

51. Jo, S., Lim, J. B., Klauda, J. B. & Im, W. CHARMM-GUI Membrane Builder for Mixed Bilayers and Its Application to Yeast Membranes. Biophys J 97, 50–58 (2009).

52. Hoover, W. G. Canonical dynamics: Equilibrium phase-space distributions. Phys Rev A 31, 1695–1697 (1985).

53. Nosé, S. A molecular dynamics method for simulations in the canonical ensemble. Mol Phys 52, 255–268 (1984).

54. Berendsen, H. J. C., Postma, J. P. M., Gunsteren, W. F. van, DiNola, A. & Haak, J. R. Molecular dynamics with coupling to an external bath. J Chem Phys 81, 3684–3690 (1984).

55. Nosé, S. & Klein, M. L. Constant pressure molecular dynamics for molecular systems. Mol Phys 50, 1055–1076 (1983).

56. Parrinello, M. & Rahman, A. Polymorphic transitions in single crystals: A new molecular dynamics method. J Appl Phys 52, 7182–7190 (1981).

57. Darden, T., York, D. & Pedersen, L. Particle mesh Ewald: An N ·log(N) method for Ewald sums in large systems. J Chem Phys 98, 10089–10092 (1993).

58. Hess, B., Bekker, H., Berendsen, H. J. C. & Fraaije, J. G. E. M. LINCS: A linear constraint solver for molecular simulations. J Comput Chem 18, 1463–1472 (1997).

59. Smart, O. S., Neduvelil, J. G., Wang, X., Wallace, B. A. & Sansom, M. S. HOLE: a program for the analysis of the pore dimensions of ion channel structural models. Journal of molecular graphics 14, 354-60–376 (1996).

60. Michaud-Agrawal, N., Denning, E. J., Woolf, T. B. & Beckstein, O. MDAnalysis: A toolkit for the analysis of molecular dynamics simulations. J Comput Chem 32, 2319–2327 (2011).

61. Humphrey, W., Dalke, A. & Schulten, K. VMD: Visual molecular dynamics. J Mol Graphics 14, 33–38 (1996).

